# Ciprofloxacin Resistance in Hospital and Community-acquired Urinary Tract Infections by Escherichia coli: A Systematic Review and Meta-analysis

**DOI:** 10.1101/2020.04.09.034041

**Authors:** Guanyu Zhou, Xiaoju Lv

**Affiliations:** Center of Infectious Disease, Sichuan University West China Hospital, China

**Author notes:** Corresponding author: Xiaoju Lv, M.D., Center of Infectious Disease, Sichuan University West China Hospital, Chengdu, Sichuan, China.

**Keywords:** Urinary tract infections, ciprofloxacin resistance, E. coli

## Abstract

In recent years, antimicrobial resistance has been increasingly reported. One main concern is the resistance of gram-negative bacteria like E. coli to ciprofloxacin (fluoroquinolones). Gram-negative bacteria are the main cause of community and hospital-acquired urinary tract infections (UTI). We aimed to review and analyze the data on ciprofloxacin resistance in hospital and community-acquired UTI. A literature search of three electronic databases (PubMed, Medline, and Cochrane) was performed. We considered the papers that were published from January 2004 to May 2019. The search yielded a total of 16097 studies besides 31 studies from a manual search. Filtering yielded 1297 relevant full-text papers. Eighty-three papers, equivalent of 99 cohorts, were finally included in this systematic review and in the analysis. The analysis results suggest that pooled ciprofloxacin resistance for community and hospital-acquired E. coli UTI is 0.27 (95% CI 0.246–0.303) and 0.30 (95% CI 0.22–0.38), respectively. Pooled resistance rates according to regions are 0.43 (95% CI 0.31–0.54) for Asia ensued by Africa 0.31 (95% CI 0.22–0.35), the Middle East 0.21(95% CI 0.13-0.30), Europe 0.18 (95% CI 0.13-0.22), and Australia 0.06 (95% CI 0.04-0.08). The pooled estimates revealed that ciprofloxacin resistance was higher in developing countries compared to that in developed countries, 0.35 (95% CI 0.30-0.40) and 0.13 (95% CI 0.10-0.16), respectively. Finally, plotting resistance over time deemed statistically significant (*n*= 79, r= 0.29, *p*= 0.038). Our findings suggest that ciprofloxacin resistance among UTI patients is a highly prevalent and serious issue. The suggested risks are low-income, acquiring hospital infection, and falling in highly-vulnerable regions like Asia and Africa. We also shed light on some approaches to correct the perception of patients and general practitioners (GPs) for antibiotic usage. We also suggest ideas to impede the progress of the post-antibiotic era in countries known for high antibiotic resistance.

## Introduction

UTI (Urinary Tract Infection) is inflammation of any tissue from the renal cortex down to the urethral meatus ensuing pathogenic invasion. It results from gram-positive, gram-negative, or candida infection (1, 2); gram-negative E. coli has been the most common cause followed by Klebsiella pneumonia (3, 4). Pseudomonas aeruginosa, staphylococcus spp., and Proteus spp. are other causative organisms (5). UTIs occur in either nosocomial/community-acquired (CA) or Hospital-acquired (HA) forms. HA-UTI is the infection that occurs: (a) after 48 hours of hospital admission, (b) before 48 hours but after invasive procedures like receiving intravenous medications (6), (c) within 90 days from two-day- or-more admission in an emergency hospital, or (d) within 30 days after undergoing invasive urinary procedures such as urinary catheters (7). It is worth noting that each day with a urinary catheter carries a 3%–7% increased risk for acquiring UTI (8). On the other hand, CA-UTI depicts infections contracted from the community or contracted during the first 48 hours of hospital admissions in the absence of any invasive interventions.

Empirical antibiotics such as fosfomycin, fluoroquinolones, β lactams, Nitrofurantoin, and pivmecillinam used to be the mainstay of management for UTI cases (9). However, this random approach increased the recurrence rates and resulted in the emergence of resistant, multi-resistant, and even pan-resistant strains. Resistance can be due to genetic mutations that, not only help the organisms survive but also pass to subsequent generations. Other explanations for resistance are anti-microbial overprescription (10) and non-prescription purchases (11). The recently-reported resistance rates have limited many therapeutic choices and underscored the significance of performing urine cultures before any empirical treatments.

Ciprofloxacin is a fluoroquinolone drug. It blocks topoisomerase type II subtype β (TOP2β), the enzyme that streamlines the supercoiling process of the mitochondrial DNA (mtDNA). Blocking this enzyme accumulates the supercoiled DNA and impedes organism replication as a result (12). Ciprofloxacin has been the most effective empirical treatment for UTI, but patients are becoming more resistant to it. For example, the resistance raised from 24.9% in 2010 to 40.7% in 2015 among Brazilian patients treated for E.coli UTI (13). Herein, we review and update rates of ciprofloxacin resistance and compare resistance in CA-UTI with that in HA-UTI caused by gram-negative E. coli.

## Methodology

### Search strategy

A comprehensive review of the databases PubMed, Medline, and Cochrane databases was performed to get the articles reporting ciprofloxacin resistance in UTI caused by E. coli. The search was run using the medical subject headings “resistance”, “urinary tract infection”, and “Escherichia coli”, and it included papers from January the 1^st^ 2004 to May the 24^th^ 2019. Reporting of this review conformed to the Preferred Reporting Items for Systematic Reviews and Meta-Analyses (PRISMA) (14).

### Criteria for Inclusion and exclusion

Papers with data on prevalence or incidence of ciprofloxacin resistance in CA or HA E.coli UTI were included. The main inclusion criteria were: (1) English articles published between 2004 and 2019 in peer-reviewed journals, (2) studies conducted on either adults or children. Nonrelevant papers with the following characteristics were excluded: (1) grey literature, (2) comments, (3) letters to editors, (4) conference abstracts, (5) reports, (6) theses, (7) non-peer reviewed studies, (8) review articles, and (9) articles with samples unrepresentative of the general population. For example, studies conducted on compromised patients or patients hospitalized for reasons other than UTI like diabetes mellitus, geriatric patients, renal transplant patients, intellectual disability, cancer, etc. Also, the authors excluded the papers nonconformist to the CDC diagnostic criteria. The CDC criteria necessitate detecting ≥10^5^ CFC/ml on urine culture (15). However, Fasugba et al. (16) was the only exception since it included samples only when more than 10^7^ CFC/ml. sparse infection (<10^5^ CFC/ml in association with UTI symptoms) and that did not specify the UTI subtype. Finally, the authors excepted papers that reported sensitivity or susceptibility with no clear resistance rate; combined or multiple resistance; and ciprofloxacin resistance in extended-spectrum beta-lactamase-resistant E. coli.

### Methods of review

Two reviewers have independently screened the titles and abstracts of the potentially-included papers. The full-texts of the papers conforming to the inclusion criteria were again screened by two independent reviewers to prove or disprove relevance.

### Data extraction

Two authors have independently extracted all the required data from the finally included articles. They settled conflicts through discussion. They developed an excel sheet to collate the following information: (1) author name, (2) year, (3) study design, (4) risk of bias, (5) study setting (community or hospital), (6) age group (adults and children or adults only), (7) country, (8) economic standard, and (9) region (Africa, Asia, Middle East, Europe, and North and South American). Specific data included the duration of each study in months, the number of E. coli positive urine samples, and the number of ciprofloxacin-resistant samples. All data are tabulated in **Table 1**.

**Table 1.**
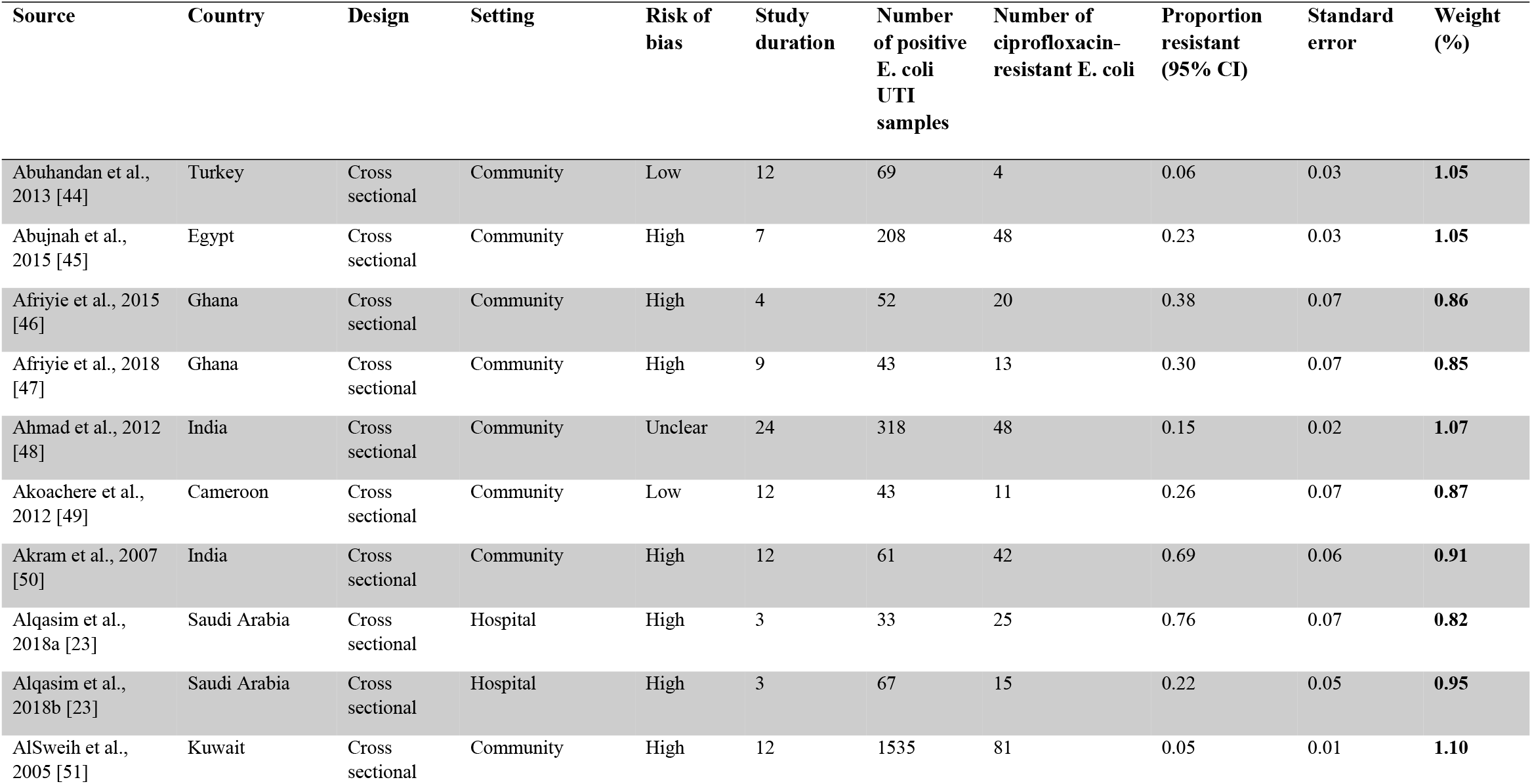
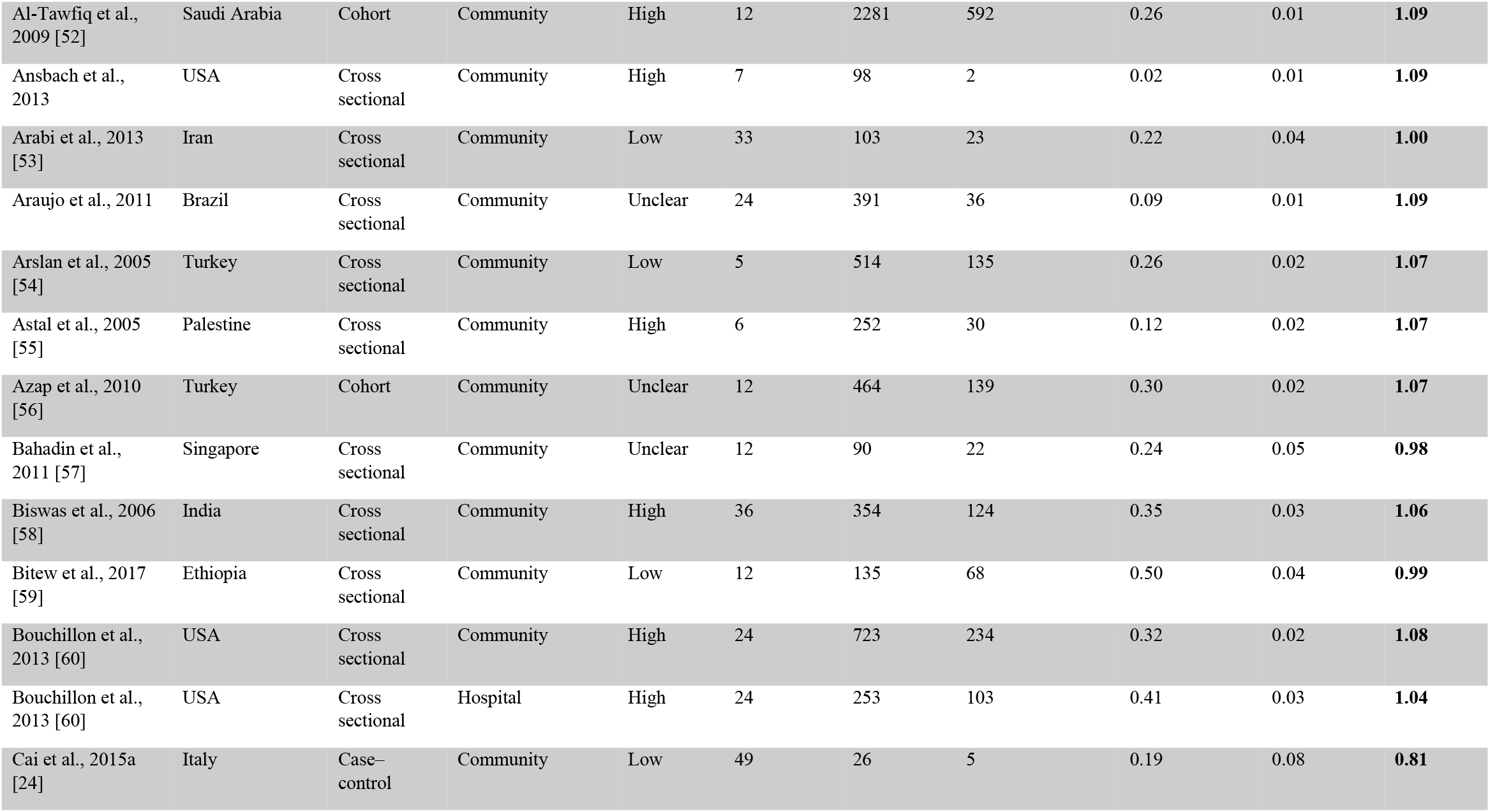
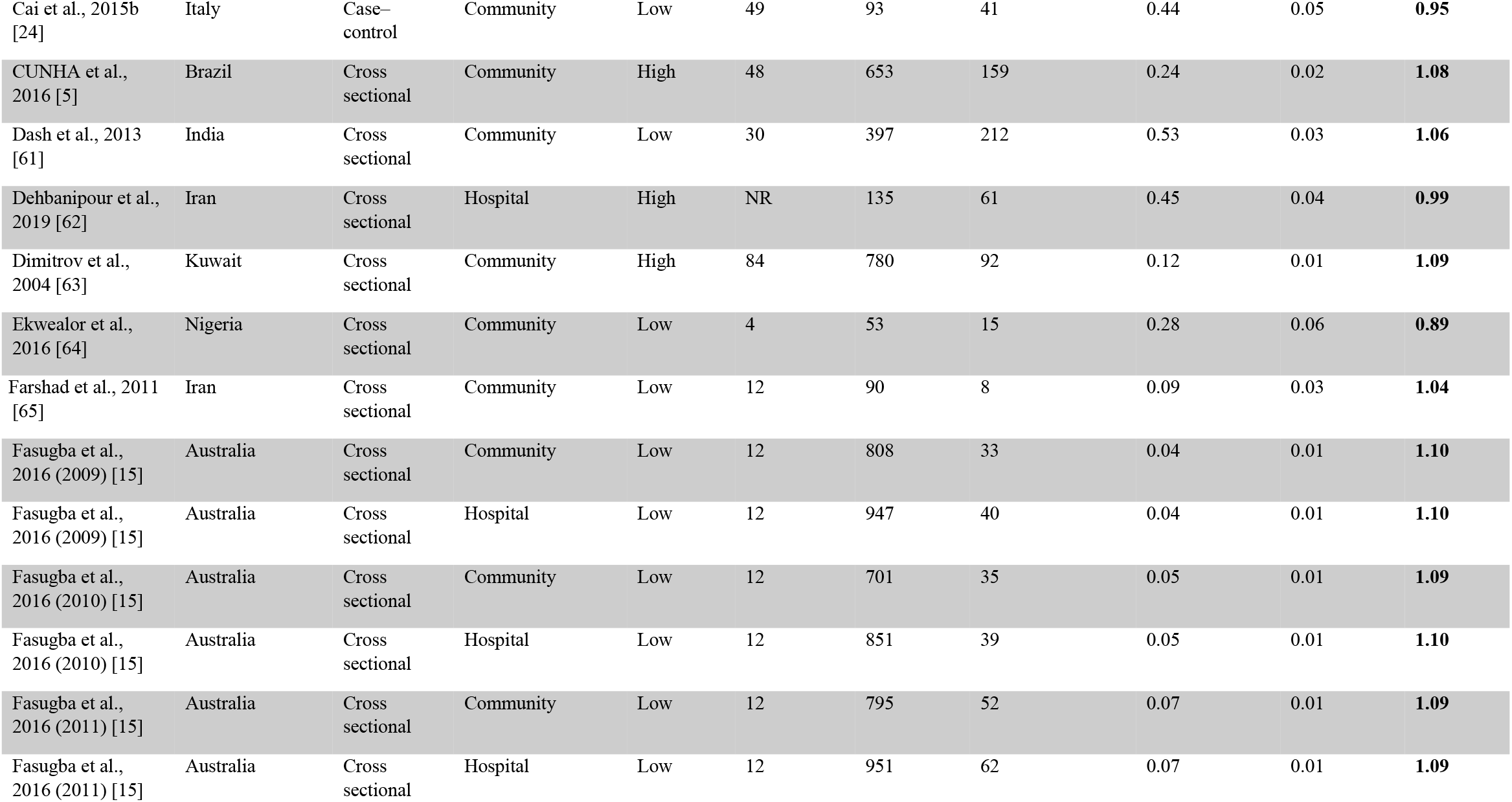
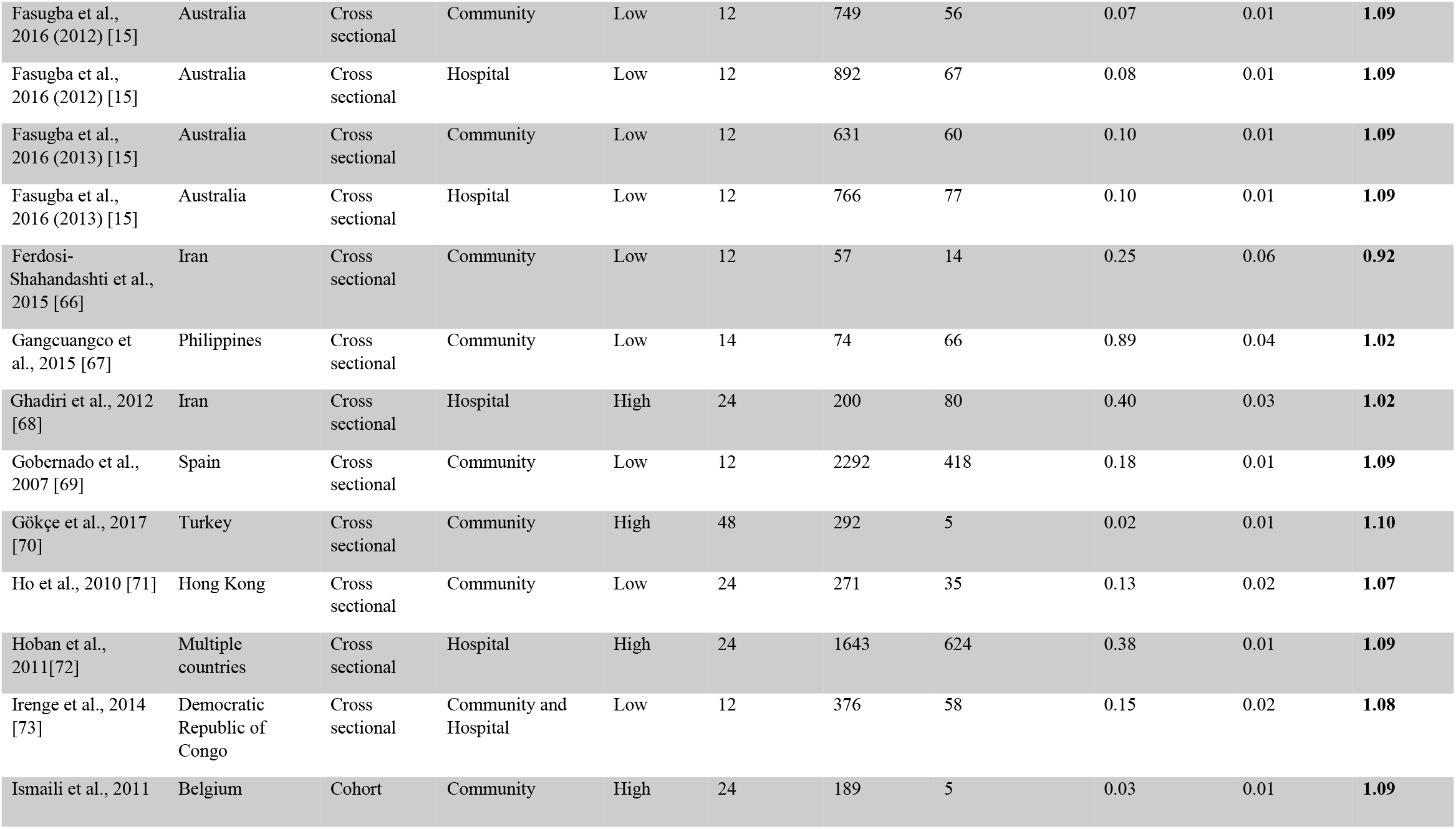
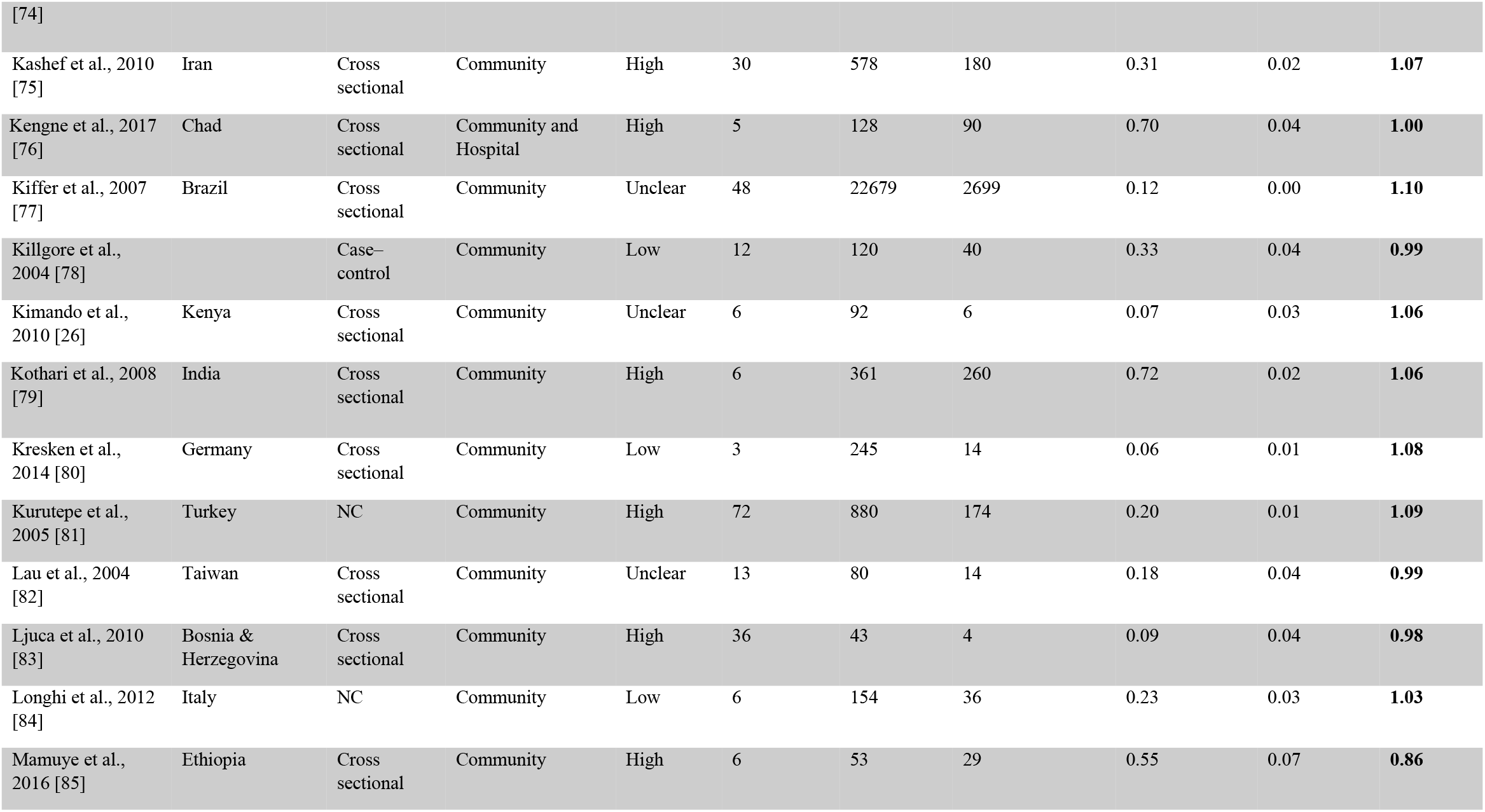
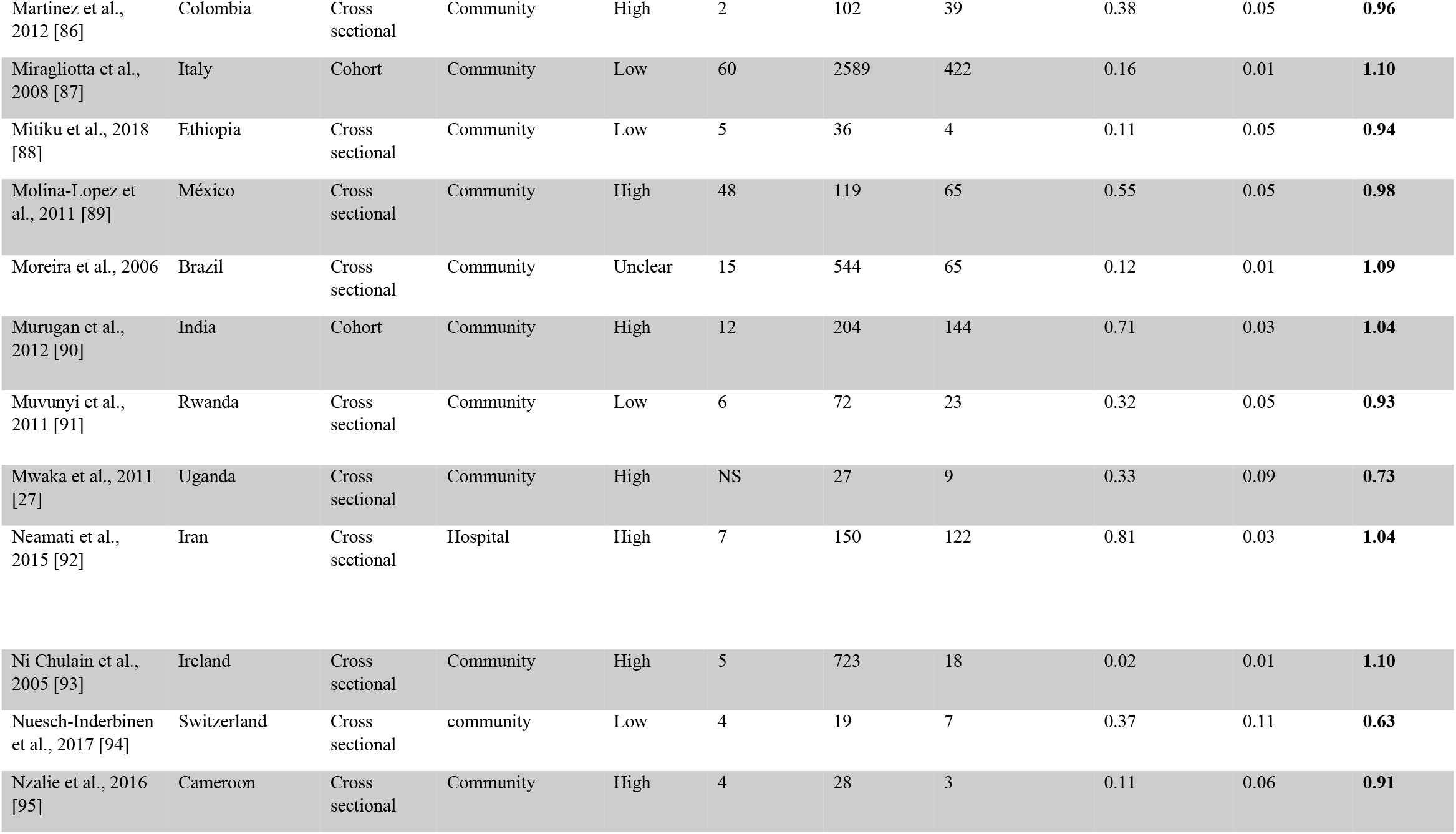
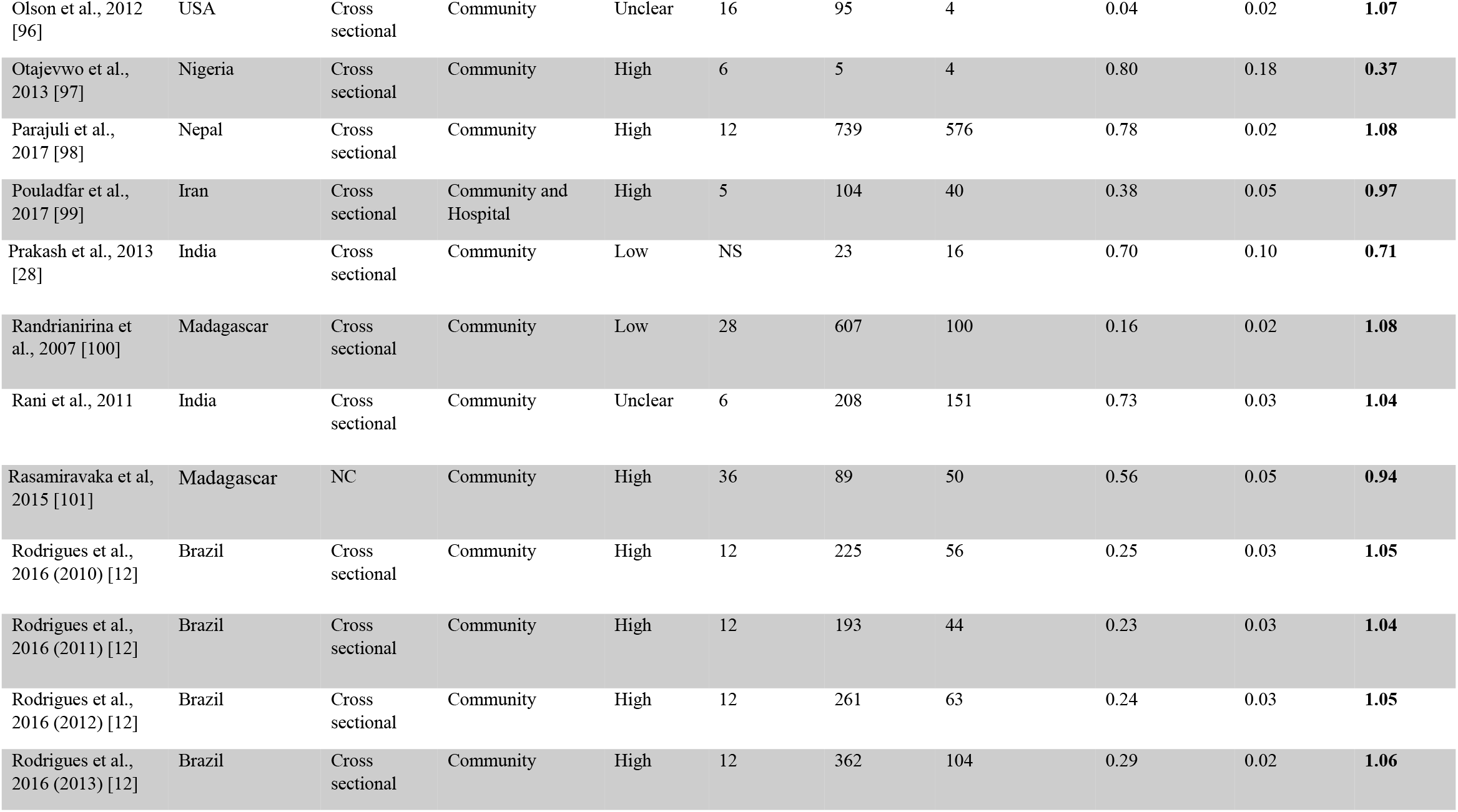
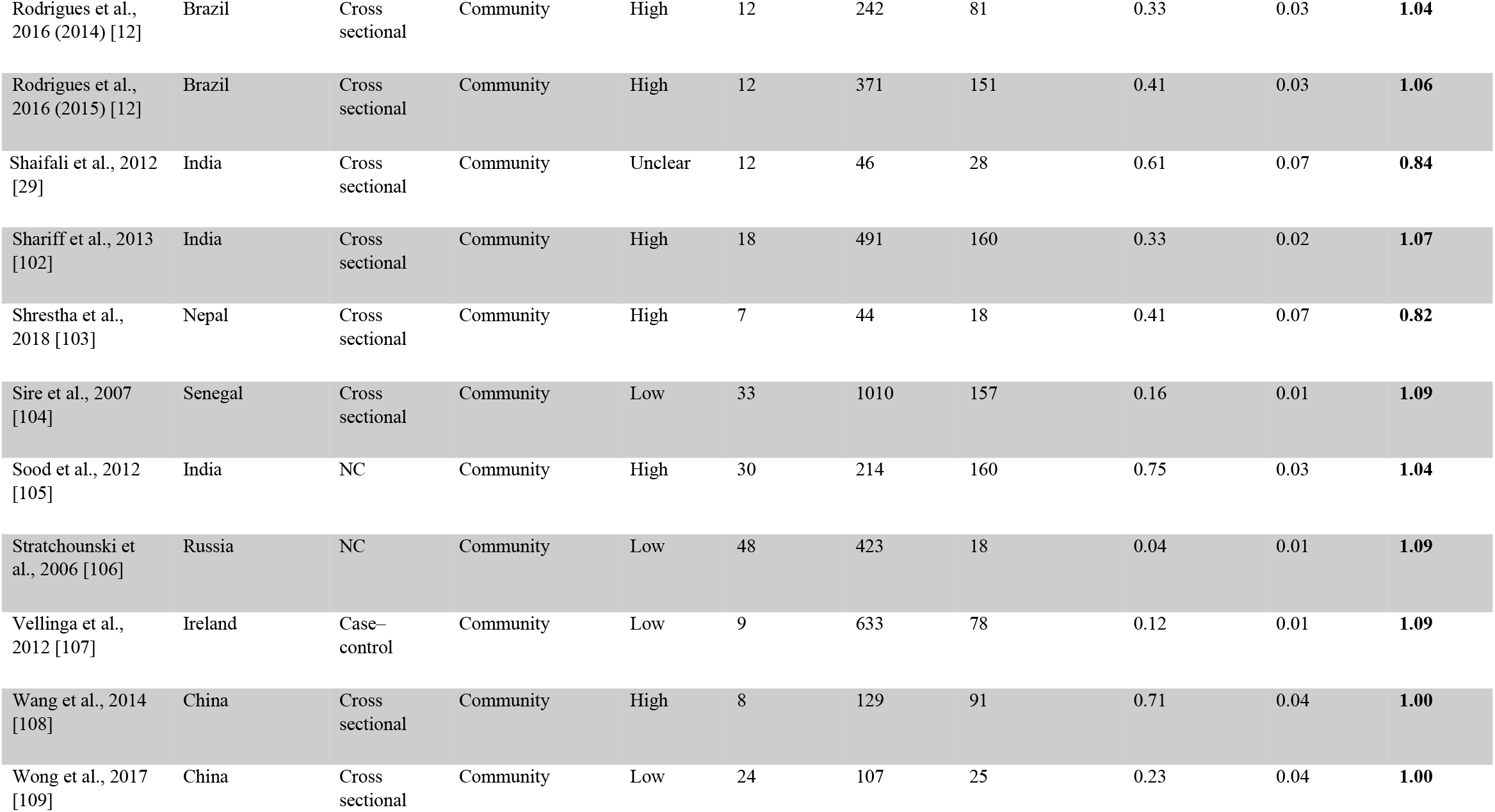
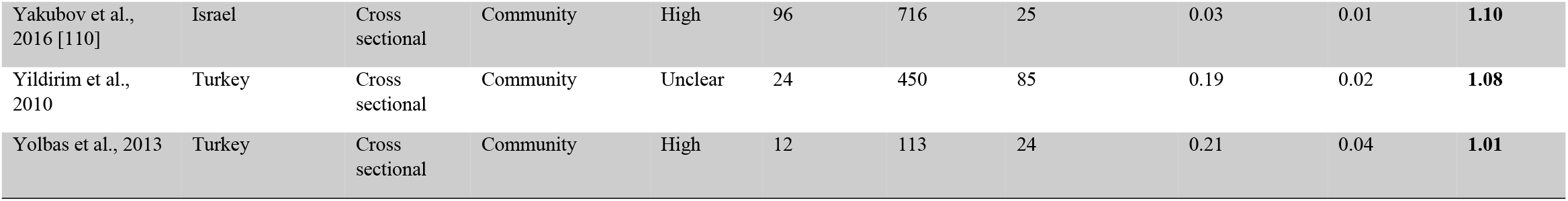
Baseline Characteristics of the Included Studies.

### Quality Assessment

The quality of the included studies was assessed using the Newcastle-Ottawa Quality Assessment Scale (NOS)(17) which classifies the quality of the analyzed studies as low, unclear, or high.

### Statistical Methods

Statistical analysis of the collated data was performed using OpenMeta Analyst software and STATA statistical software version 14 (18). The authors calculated the pooled proportions with 95% confidence intervals for ciprofloxacin resistance in patients with E. coli UTI; they reported them both separately and as a comparison between the community and hospital settings. The random-effects model (DerSimonian and Laird’s method) (19, 20) was used to evaluate heterogeneity among the studies. Statistical heterogeneity was reported using the chi-square-based Q and the I^2^ statistic (21). Sensitivity analyses were conducted to assess the heterogeneity and robustness of the pooled results; sensitivity results deemed statistically significance at P-value<0.05. The authors did subgroup analyses to investigate ciprofloxacin resistance for each potential risk factor: the risk of bias, economy, region, and population. Meta-regression analysis (22) was conducted to examine the observed heterogeneity and to detect the potential factors beyond. A funnel plot estimated the publication bias. Also, Spearman’s correlation coefficient with the median study year reported the significance of ciprofloxacin resistance over time. For studies occurred over two years, the first year was analyzed, and for the papers occurred over four years, the second year was analyzed. Finally, if records occurred over six years, the third year was analyzed.

## Results

### Search results

Electronic searches detected 16097 papers and 31 more papers through manual searching. After removing 11366 duplicates, 4731 papers were eligible for the following stage of title and abstract screening. The screening qualified 1297 papers for full-text screening. After scrutinizing the inclusion and exclusion criteria, 83 papers were relevant. These papers displayed data from 99 cohorts varying between HA and CA-UTI. The discrepancy between numbers is due to multiple sampling. For example, Rodrigues et al., 2016 reported resistance in CA-UTI annually and consecutively from 2010 to 2015 (13); accordingly, their study was considered as six papers. Similarly, Fasugba et al., 2016 detected the resistance in both CA and HA cases over five years; they reported ten cohorts (16). Alqasim et al., 2018 (two cohorts) compared the ciprofloxacin resistance in two E. coli strains (23). And so were Cia et al., 2015 who compared two groups of untreated and previously treated patients (24). **Figure 1** displays the PRISMA flowchart for the search steps and exclusion reasoning.

**Figure 1.**
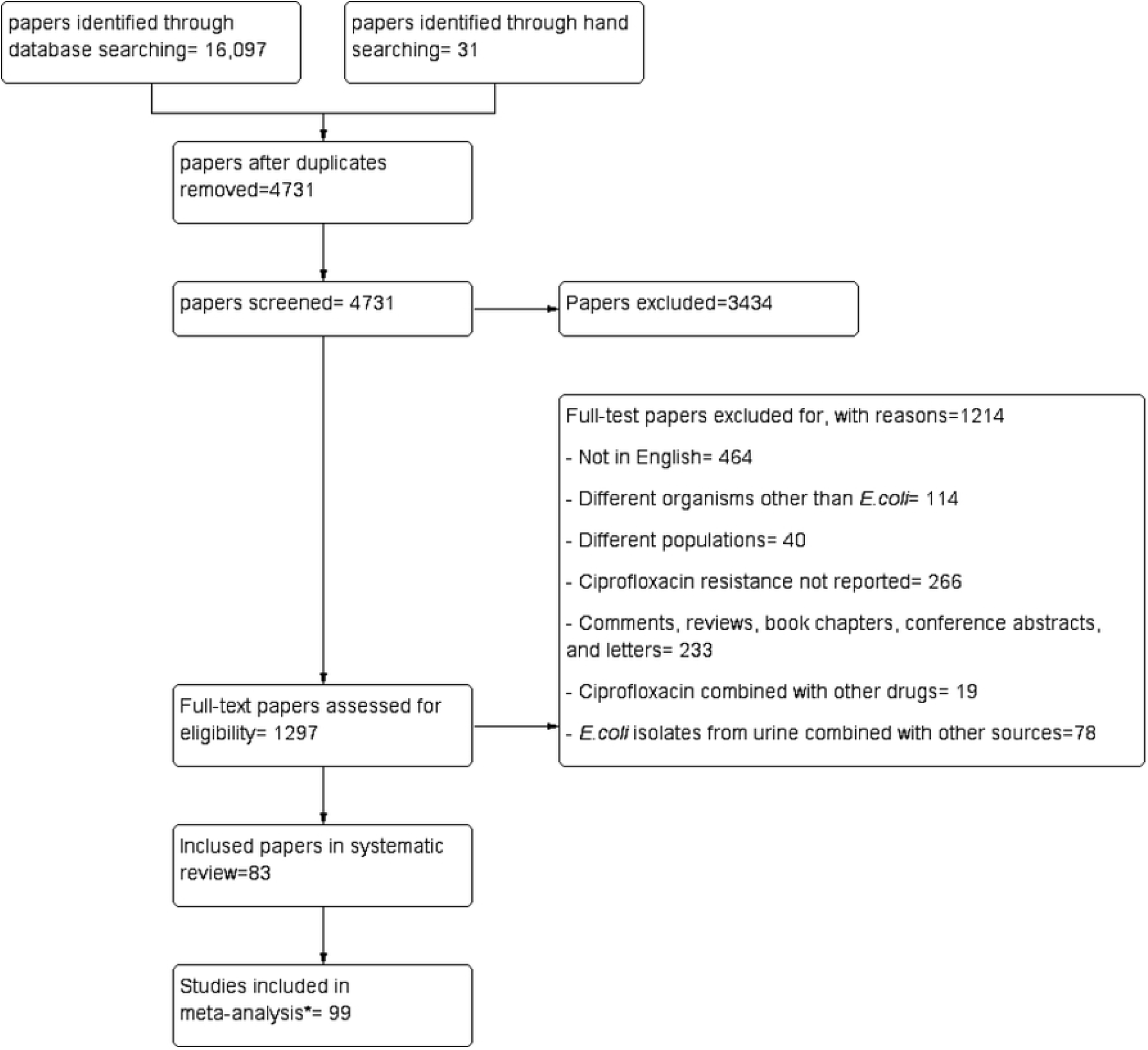
PRISMA flow chart. (*99 reports from 83 papers)

### Study characteristics

Out of 83 studies investigating ciprofloxacin resistance, 28 (34%) studies were in developed countries, and 55 (66%) were in developing countries. Geographically, 17 (20%) studies were in Africa, 12 (14%) studies in the Americas, 25 (30%) in Asia, one (1%) in Australia, 17 (20%) in Europe, 11 (13%) in the Middle East, two (2%) in South America, and one (1%) in multiple countries. The duration ranged from two to 96 months, and the majority of the studies (88%; n=73) detected resistance in patients in community settings. Data on age and sex of E. coli UTI patients subjects were reported in 93% (n= 77) and 96% (n= 80) of studies, respectively. Table 1 presents further detail on the characteristics of the included studies.

### Risk of bias

The methodological quality of the included papers was assessed using the Newcastle Ottawa Scale from Cochran collaboration (17); it revealed 49% (n= 44) high-risk of bias articles, 34% (n= 28) low-risk of bias articles, and 13% (n= 11) unclear-risk of bias articles.

### Pooled ciprofloxacin resistance (setting)

The pooled estimate for ciprofloxacin resistance in CA E. coli UTI was 0.27 (95% CI 0.246-0.303, **Figure 2**) as opposed to that in the hospital setting 0.30 (95% CI 0.22-0.38, **Figure 3**). There was substantial heterogeneity between community and hospital studies (I^2^= 98.95, p <0.001 and I^2^= 99.28, p <0.001, respectively). However, ciprofloxacin resistance in HA E. coli UTI patients was significantly higher than its rival (p <0.001). **Figure 4** shows a forest plot of the studies reporting ciprofloxacin resistance in CA-UTI patients by region. The plot revealed that Asia has the highest pooled ciprofloxacin resistance 0.43 (95% CI 0.31-0.54) ensued by Africa 0.31 (95% CI 0.25-0.30), the Middle East 0.21 (95% CI 0.13-0.30), Europe 0.16 (95% CI 0.12–0.21), and finally Australia 0.06 (95% CI 0.04-0.08). Countries were classified as developed and developing according to the World Bank classification of 2018 (25), another pooled estimate was measured predicated on the income. The analysis showed a significantly higher pooled ciprofloxacin resistance (p < 0.001) in developing countries compared to developed countries. 35 (95% CI 0.30-0.40) and 0.13 (95% CI 0.10-0.16), respectively **(Figure 5)**.

**Figure 2.**
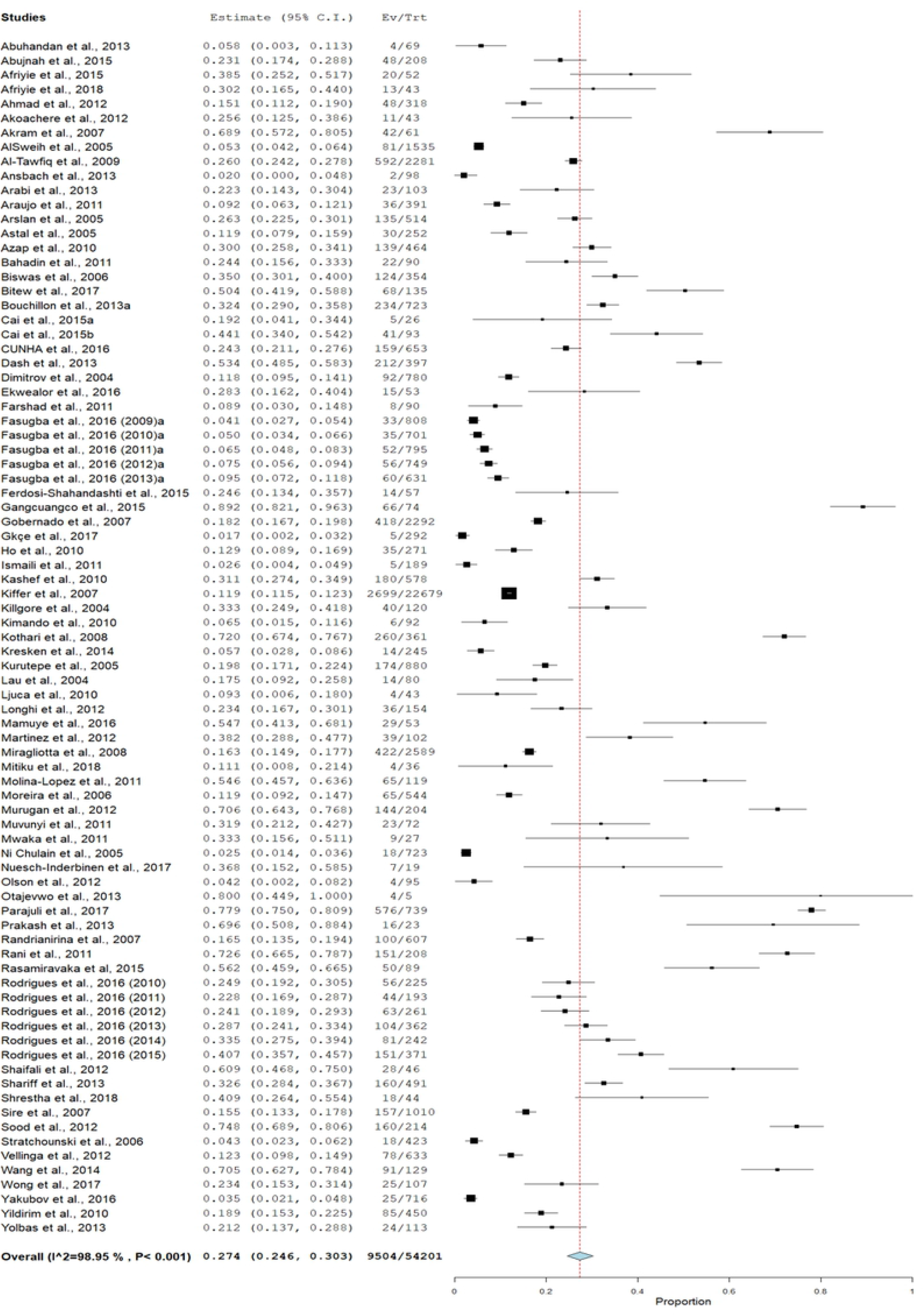
Forest plot of Ciprofloxacin resistance in community-acquired E.coli UTI.

**Figure 3.**
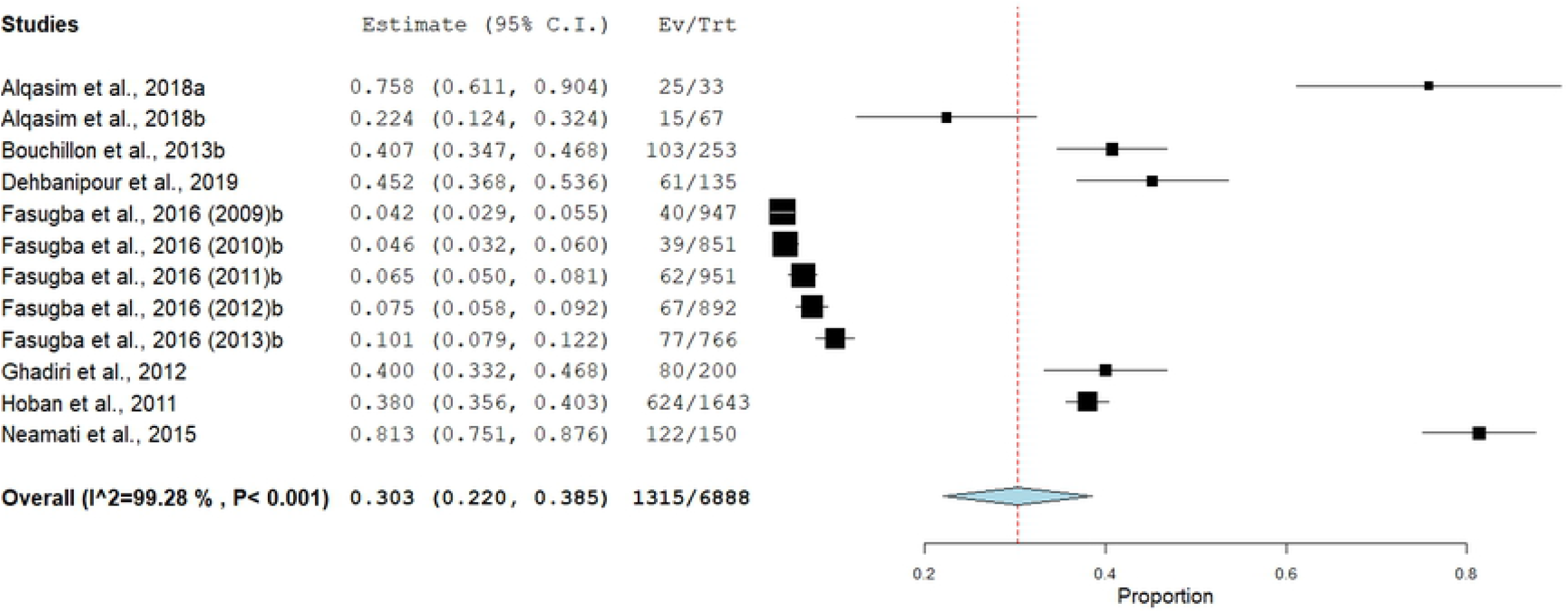
Forest plot of Ciprofloxacin resistance in hospital-acquired E.coli UTI.

**Figure 4.**
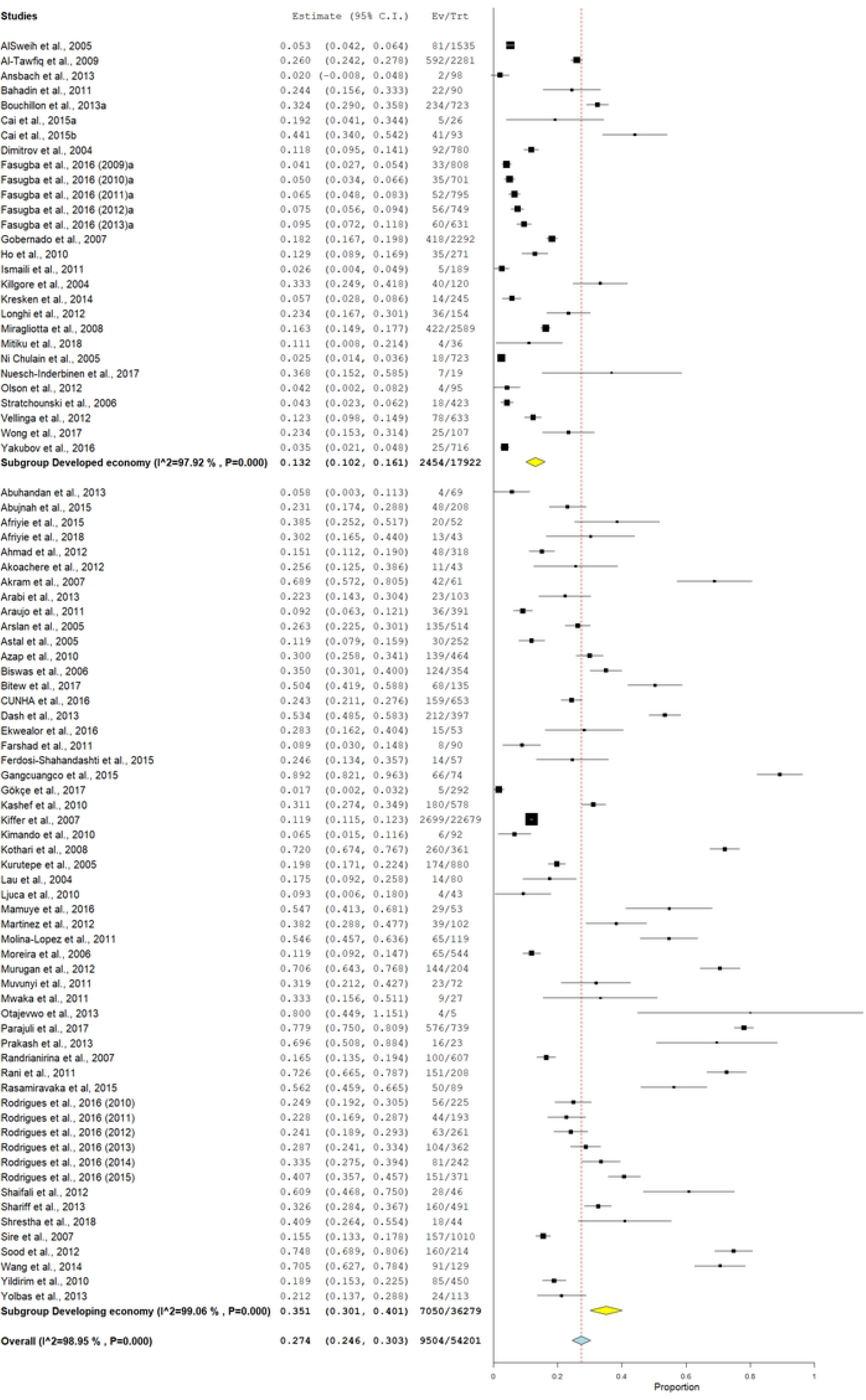
Forest plot of Ciprofloxacin resistance in community-acquired E.coli UTI by economy (developed or developing country).

**Figure 5.**
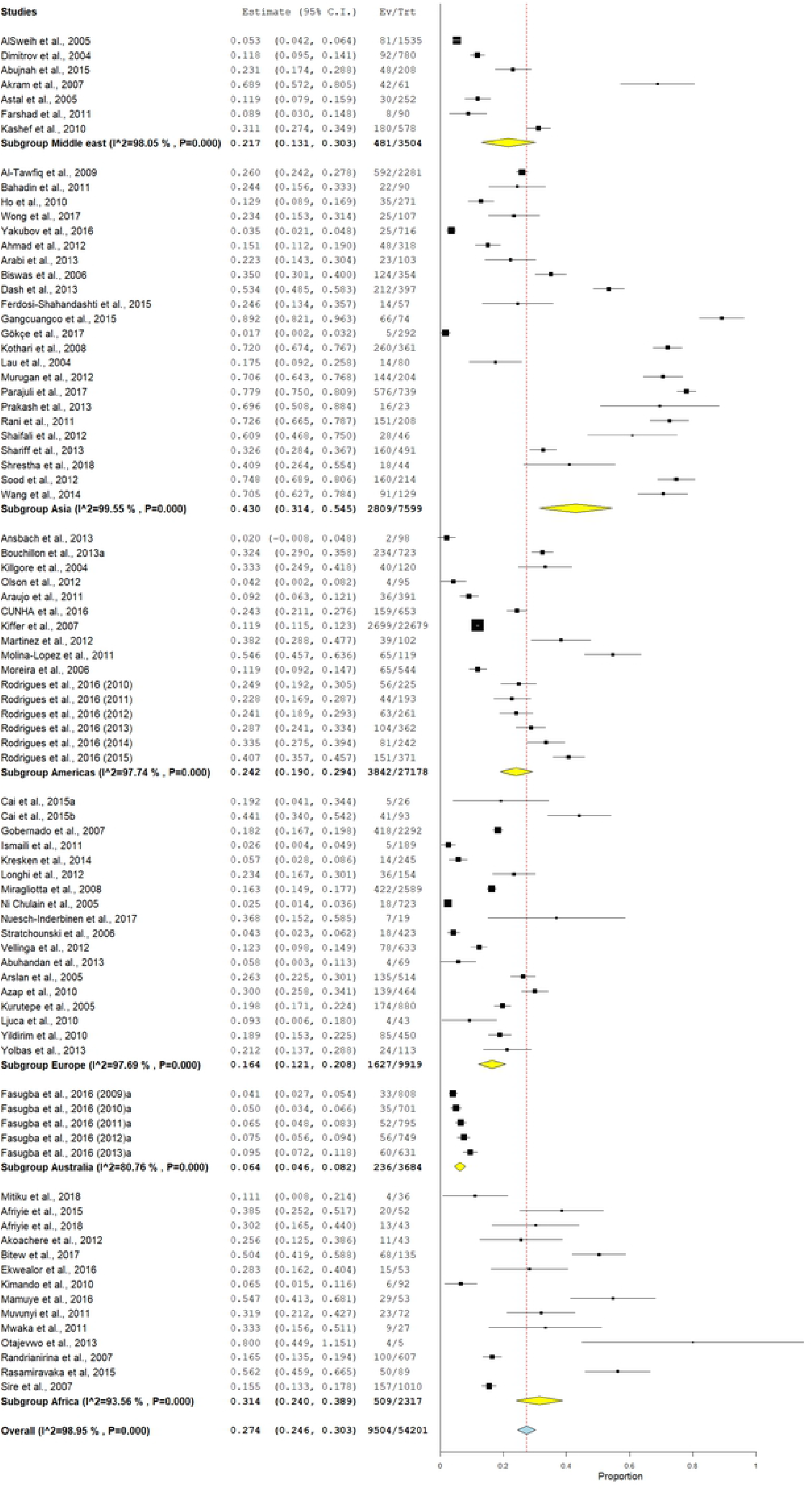
Forest plot of Ciprofloxacin resistance in community-acquired E.coli UTI by region.

### Resistance over time

Only four studies did not report the year when they were conducted (26–29). Data from 79 studies were plotted to depict the changes in ciprofloxacin resistance over the years (**Figure 6**). Spearman’s correlation coefficient showed a positive yet weak relationship between the development of resistance and the elapse of years; it also reflected a statistically significant rise in ciprofloxacin resistance with time *(n=* 79, r= 0.29, p= 0.038). Further analysis revealed a significant correlation in developing countries. Conversely, there was no significant increase in ciprofloxacin resistance over time in developed counties.

**Figure 6.**
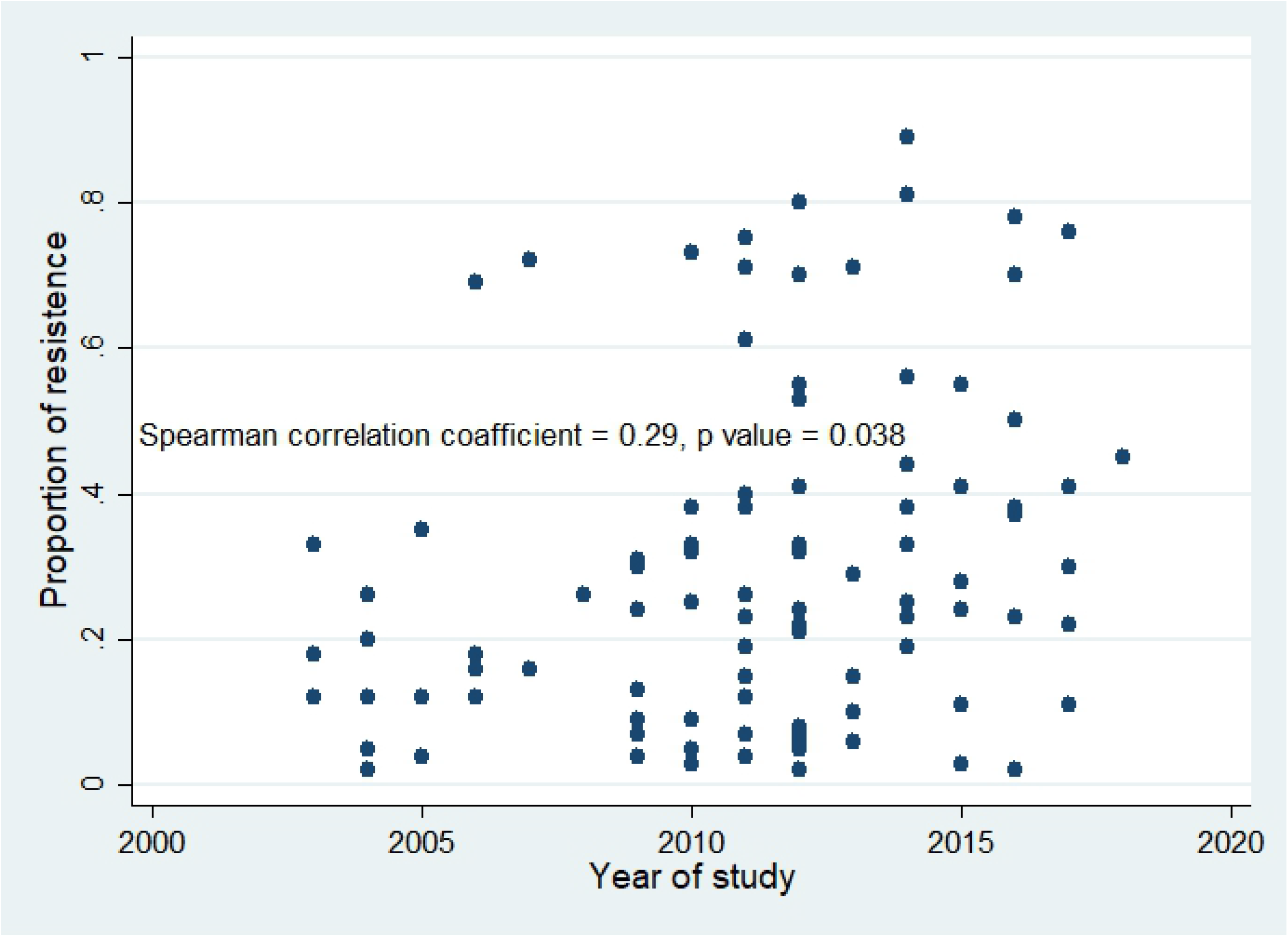
Scatter plot of ciprofloxacin resistance in community-acquired UTI over time; N = 79 (4 studies excluded due to missing information on year study was conducted).

### Subgroup analysis

Subgroup analysis of community infection detected a significant difference in pooled ciprofloxacin resistance as per risk of bias, economy, duration, age group, and region. On the other hand, there was no significant difference for both the region and economy factors in HA infections (**Table 2**). Furthermore, comparing both settings in the examined subgroups revealed significant differences (p-value <0.001) for risk of bias (high); economy (developing); and region (Americas).

**Table 2.** Subgroup analysis of ciprofloxacin resistance in community-acquired E. coli UTI patients by risk of bias, study duration, economy, region and age group.

| Subgroup | Community setting | Pooled resistance | P value |
| --- | --- | --- | --- |
|  | **N = 73** |  |  |
| Risk of bias | Low and unclear n = 39 | 0.220 | < **0.001** |
|  | High n = 34 | 0.333 |  |
| Study duration | <12 months n = 26 | 0.347 | < **0.001** |
|  | ≥12 months n = 47 | 0.251 |  |
| Economy | Developed n = 28 | 0.144 | < **0.001** |
|  | Developing n = 45 | 0.347 |  |
| Region | Africa, Asia and Middle East n = 44 | 0.364 | < **0.001** |
|  | Europe, Australia and American = 29 | 0.179 |  |
| Age group | Adults and children = 38 | 0.236 | < **0.001** |
|  | Adults = 24 | 0.310 |  |

### Meta-regression analysis

The observed heterogeneity among studies reporting CA-UTI E.coli was detected by random effects meta-regression. Factors responsible for heterogeneity were economy (0.004), Asia (0.008), and unclear risk of bias (0.021).

### Sensitivity and Publication bias

A funnel plot was generated to assess the potential publication bias based on the results of ciprofloxacin resistance proportions (**Figure 7**). The plot showed obvious asymmetry suggesting no evidence of publication bias (Egger’s p-value= 0.012). The robustness of the results was assessed by a sensitivity analysis that underscored the contribution of each study to the overall estimate in ciprofloxacin pooled proportions.

**Figure 7.**
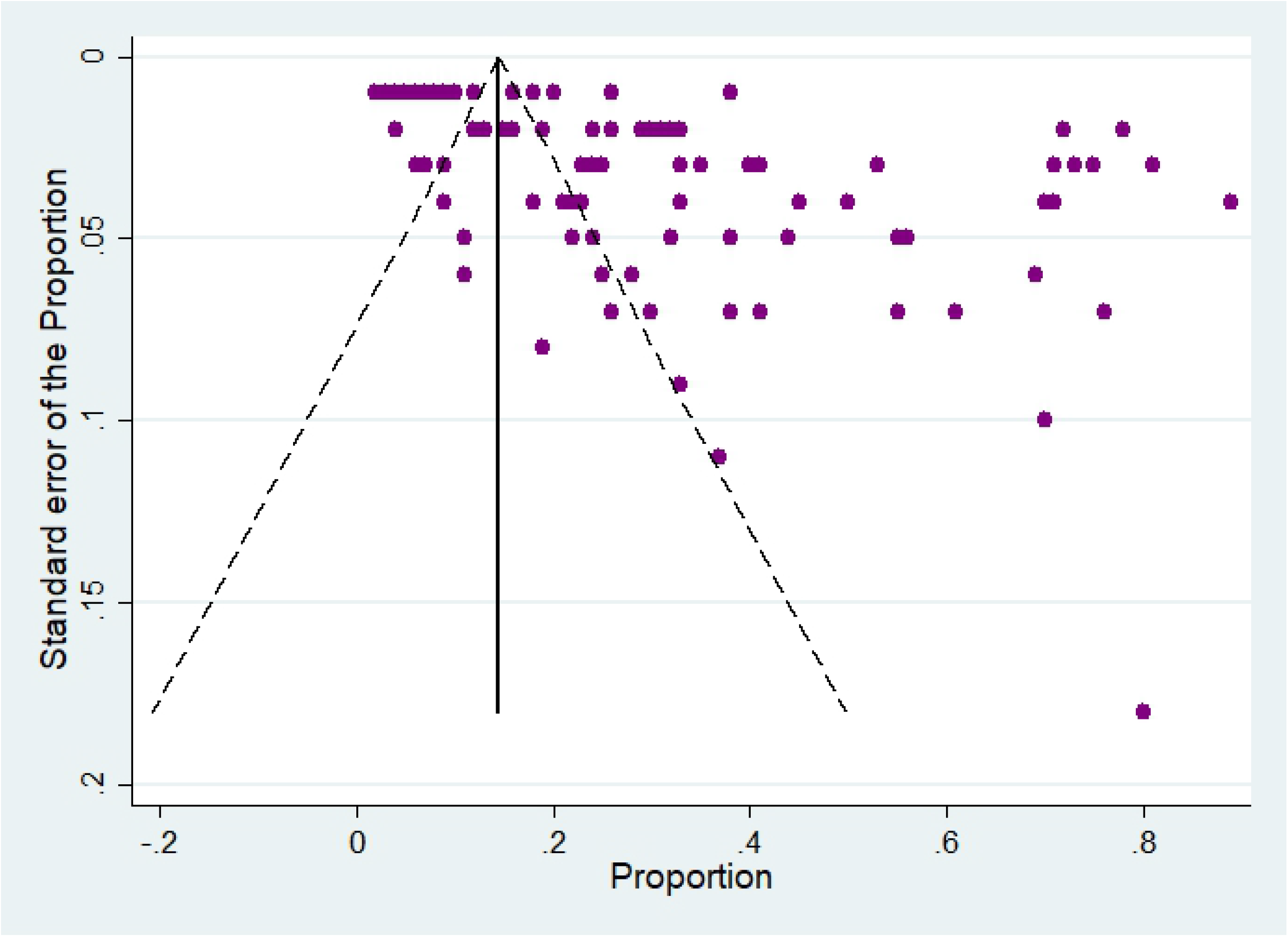
Funnel plot for the articles entailed in the meta-analysis.

## Discussion

Our review aims to compare ciprofloxacin resistance in E. coli UTI between hospital and community settings. The findings suggest a higher ciprofloxacin resistance in hospitals compared to that in the community. Furthermore, ciprofloxacin resistance was significantly higher in developing than in developed countries. Asia showed the highest resistance rate (43%), followed by Africa (31%), the Americas (20.7%), the Middle East (21.7%), Europe (16.3%), and Australia (6.4%). A similar review was conducted in 2015 (30), but the emergence of new primary studies on the same issue highlights the need for updating the data to scrutinize new evidence and compare the findings (31).

Antibiotic resistance is a growing concern. The WHO report in 2014 was the first to highlight the global magnitude of this problem and its untoward ramifications like accelerating an antibiotic era (32). One of the main reported issues was the growing resistance to fluoroquinolones in UTI caused by E. coli despite developing recently in the 1980s. Great efforts followed this report; the Who launched the GLASS (Global Antimicrobial Resistance Surveillance System) initiative in selected countries to collate data on resistance to a constellation of antimicrobials. The initial phase (2017-2018) collected data on antibiotic consumption as well, being proportionate with resistance. The data showed a growing resistance to ciprofloxacin in E. coli UTI (33).

Our findings suggest a higher resistance of E. coli UTI to ciprofloxacin in the hospital than in the community setting. This finding is comparable to Fasugba et al., 2015 (30). However, there is evidence that ciprofloxacin resistance among urology patients is specifically higher than that in CA or HA infections (34). Many reports have individually recorded skyrocketing ciprofloxacin resistance rates. For example, Zahedi et al., 2018 have reported resistance of 77.5% (35); also, El-Sokkary et al., 2015 have recorded a resistance rate of 67% in HA-UTI (36). Resistance in HA-UTI is due to contracting basically-resistant strains from hospitals. Antibiotic resistance has been explained by the vulnerability of hospital patients to infection due to poor immunities and associating comorbidities (37).

Our meta-analysis revealed higher ciprofloxacin resistance in developing countries than that in developed countries, 34.7% and 14.4% respectively. The difference was statistically significant. This finding correlates with Founou et al., 2017 who, additionally, touted antibiotic resistance as responsible for 90% of mortality from infections in 11 developing countries (38); it is also comparable to results of Fasugba et al., 2015 (30). What increases the credibility of these comparisons is that both studies classified countries according to the World Bank Classification. Poverty and lack of knowledge are the main contributors to developing antibiotic resistance in developing countries. These factors increase the over-the-counter and empirical use of antibiotics. Patient expectations are another player; 40% of GPs (General Practitioners) prescribe antibiotics only to meet the expectations of their patients (39). Besides growing resistance, developing countries should be vigilant of the correlation between antibiotic resistance and out-of-pocket expenditure; the latter increases the poverty rate in a vicious circle (40). These data highlight the importance of launching campaigns to educate the public on the appropriate use of antibiotics. Moreover, governments of the developing regions should implement interventions and regular surveillance programs to report the frequency of prescribing antibiotics as an initial step in measuring and containing the high resistance rates.

Further analysis based on regions revealed that Asia (43%) had the highest ciprofloxacin resistance, followed by Africa (31%), the Americas (28.4%), the Middle East (21.7%), Europe (16.3%), and finally Australia (6.4%). Economic risk supports this regional difference; for example, Asia has many low and middle-income (developing) countries. This finding is in line with that of Jean et al.; they reported a fluoroquinolone resistance rate of 50% and added that fluoroquinolones are no longer apt for treating UTI in the Asia-Pacific region. However, they did not mention the setting or sample size calculations (41). That is also comparable to Fasugba et al., 2015 who even reported higher resistance (50% versus 43%) (30). Decreased resistance may reflect modest efforts to reduce resistance in this region since 2015.

It is central to emulate the antibiotic consumption plans followed by countries like Australia (resistance rate of 6.4%). In a study investigating knowledge, attitude, and behavior of the Australian GPs towards prescribing antibiotics, Gps deemed prudent when using the available medications. They stick to symptomatic treatment and do not rush to antibiotics (39). Moreover, Australia adopts approaches to minimize the use of antibiotics. Australian GPs are encouraged to share decisions with their patients and to delay and not to repeat antibiotic prescription. Also, the Australian government has implemented a ban on some antibiotics like fluoroquinolones (ciprofloxacin, norfloxacin, and moxifloxacin) (42). Accordingly, ciprofloxacin is preserved only for serious gram-negative infections resistant to other antibiotics; this practice explains our finding of low resistance to ciprofloxacin in Australia. Cheng et al., 2012 reported similar results, yet their main objective was proving the efficacy of the approach aforementioned in precluding the progress of quinolone-resistant strains. They reported a rise in resistance from 1% in 1998 to 5.2% in 2010. This resistance skyrocketed across the United States from 3% in 2000 to 17% in 2010, the year when resistance to quinolones across Europe was 45% (42). Of note, some European countries (the second only to Australia in low ciprofloxacin resistance) like Sweden, despite not being represented in our analysis, has held record for eliminating unnecessary prescriptions and reducing resistance rates over the past 20 years. More research on similar countries is needed to recap methods for implementing such evidence-based approaches.

Additionally, our meta-analysis of ciprofloxacin resistance over the years from 2004 to 2019 has revealed a positive and statistically significant yet weak correlation (n= 79, rs= 0.29, p= 0.038). Fasugba et al., 2015 reported a more statistically significant and relatively stronger correlation (n= 47, rs = 0.431, p=0.003) (30). Low significance in our article is possibly due to the large number of included papers and the potential missing data.

### Strengths and limitations

To our knowledge, this review is the largest to investigate ciprofloxacin resistance in E. coli UTI (CA and HA-UTI). We included a large number of cohorts (99 cohorts from 83 articles). Moreover, this has been the first review articles since 2015, which means that there is plenty of studies and power to represent geographically large areas and to detect effects. We specified the settings and patient criteria to ensure the ecological validity of the included studies, so our findings are applicable for these settings. Although the number of studies conducted in the hospital setting was fewer than that of the community setting, it was still higher than any previous study investigating HA-UTI. Other strengths are including papers only when they were published in peer-reviewed journals, only when their methodologies conformed to the CDC criteria for diagnosing UTI, and only when they mentioned the setting. The latter two have excluded many papers. However, they standardized the inclusion criteria, increased the internal validity and homogeneity of the included studies, and added to the uniformity of data. Another strength is excluding populations with specific comorbidities to boost the external validity of our findings and the reliability to generalize them. Finally, we included only papers reporting resistance. In other words, we never used intermediate susceptibility to deduce resistance rates.

We acknowledge that our review has some limitations; we noted substantial heterogeneity among the included studies and estimates. One limitation is the different diagnostic criteria among the included studies. While some papers relied on some clinical signs and symptoms in association with the laboratory findings, others relied merely on the laboratory results; this possibly included complicated cases and over valuated resistance rates. Another limitation is the lack of standard microbial susceptibility tests during screening for inclusion. Finally, the subgroup analysis by region might be uneven since some regions have exemplary countries in the realm of eliminating the use of antibiotics that were not represented in the analysis, some European regions like Sweden for example. Further research is needed to address the real resistance rates and to draw conclusions on the best ways to curb rates of antimicrobial resistance.

## Conclusions

The current review and meta-analysis revealed that some risk factors increase the resistance of E.coli strains causing UTI to ciprofloxacin. Low-income countries, hospital-acquired UTI, and falling in highly-vulnerable regions like Asia and Africa are thereof these factors. We also underscore the significance of detecting and reporting cases within the healthcare system to global sides like the WHO to handle the problem. Also, governments should increase the awareness of Gps about the proper antibiotic practices; they should adopt the approaches of the leading countries in overcoming resistance like Australia and Sweden. Finally, special attention should be geared towards the public to correct their entrenched wrong believes on antibiotics as magic therapies.

## Acknowledgment

The authors thank Shenzhen People’s Hospital for support assistance.

